# Chemical blueprints to identifying fire ants: overview on venom alkaloids

**DOI:** 10.1101/407775

**Authors:** Eduardo Gonçalves Paterson Fox

## Abstract

*Solenopsis* fire ants are remarkably difficult to identify using morphological characters, particularly from the most abundant minor workers. The present manuscript introduces a set of chemical tools to facilitate species diagnosis from field-collected fire ant samples, herein focusing on minor workers. Fire ants from different localities (native and invaded) were analysed using gas-chromatography. Samples were collected from the field into organic solvents; sampling effort included 14 species, and a suspected hybrid. A total of 32 piperidine alkaloids were spotted and tentatively identified and representative relative chemical proportions for minor workers are presented for the first time for a number of species. It is hoped that the provided info will prove useful to researchers working on fire ants in future studies. Further compounds are being analysed for additional auxiliary tools.

## Introduction

The New World fire ants comprise a group of ca. 20 species within the genus *Solenopsis*, famous for their aggressive behaviour and painful stings (Pitts et al. 2018). The fire ants constitute a monophyletic species assembly native to the Americas, and most species are known from South America (Tschinkel 2006, Pitts et al. 2018).

Fire ants are notoriously difficult to identify based on morphology alone, particularly in their native geographical range. This is mainly because of considerable plasticity and infraspecific variations in established characters, and a paucity of reliable characters applicable to the most abundant caste of minor workers (Tschinkel 2006, Pitts et al. 2018). Moreover, several species have not been clearly defined (Ross et al. 2010) and populations are known to hybridise (Tschinkel 2006, Ross et al. 2010), adding further noise to the clear delimitation of defining characters.

Using chemical characters to separate and classify fire ant species has been proposed several times in the past (e.g. Vander Meer & Lofgren 1988, DeFauw et al. 2010). Fire ants are remarkable in having conspicuous amounts apolar compounds that can be readily extracted by immersion in organic solvents (Fox et al. 2012). A number of studies have been published characterising the diversity and characteristics of such compounds (e.g. Brand et al. 1972; Chen et al. 2009; for a review refer to Chen et al. 2012). The most abundant and stable chemicals are piperidinic alkaloids which are essentially a chemical mark of fire ants (Fox 2014), aka ‘solenopsins’. Regardless of widespread intraspecific variations, the venom alkaloids of fire ants have been repeatedly demonstrated as useful in sorting between cryptic and sympatric fire ant species and populations (Dall’Aglio-Holvorcem et al. 2009; Fox et al. 2012, Araújo et al. 2018).

There are currently well defined protocols for separating and analysing fire ant venom alkaloids (Fox et al. 2013; Yu et al. 2014) as well as a number of published raw chromatograms available as chemical references (e.g. Fox et al. 2018a, Fox et al. 2018b) allowing for reproducible results.

The present manuscript presents representative patterns of venom alkaloids obtained from minor workers from different fire ant species (both nominal and cryptic) found in South America. These result from the analysis of >100 chromatograms from >30 scattered collection sites, where species identifications are supported by morphological and barcoding approaches. We hope the provided alkaloid chemistry patterns prove useful to other researchers attempting to correctly identify fire ant samples.

## Methods

### Ant samples

Fire ant nests were located during field trips, mostly by visual inspect around urban parks and highway roadsides (with the exception of *Solenopsis virulens*, which had no visible fixed nest). Workers of different castes were collected by the numbers 3-5 into either into hexane or ethanol in glass vials using fine tweezers, according with local conditions. Only minor workers are herein presented, given their venom alkaloids patterns are more stable across species (EGPF unpublished results), and the fact that they are the most abundant caste in a fire ant nest (Tschinkel 2006). Species identification was based on morphological analysis after Pitts et al. (2005), mainly from accompanying major workers characters. Auxiliary molecular mtDNA identifications were kindly provided by Dietrich Gotzek, based on Ahrens et al. (2009). Voucher specimens deposited at Instituto Biologico Sao Paulo, MNRJ / UFRJ Brasil, HSK University China.

### Chromatograms

Given circumstances the gas-chromatographs machinery used included different brands (see Acknowledgments for details) but always set to the same settings (detailed Fox et al. 2018) enabling reproduction of easily compatible results patterns. Settings in short were: injection volume 1 μL; RTX-5MS silica capillary column (30 m × 0.25 mm × 0.25 μm); carrier gas helium at flow rate 1.0 mL/min; MS scanned at 70 eV set to speed 1228. Temperatures: injection port 250°C; oven temperature increasing from 50°C to 290°C at 12°C/min followed by a paused hold of 6 minutes and a final 30°C/min increase from 290°C to 320°C to clean column from residues.

Samples of every caste of the well-studied, alkaloid-diverse fire ant species *Solenopsis invicta* Buren (e.g. see Tschinkel 2006) were analysed with each different machine ensuring references in case of ambiguous peak identifications. Further ambiguous identifications were resolved by comparing with hydrocarbons, whenever available, both as internal and as external standards. The species-specific cuticular hydrocarbons patterns are more complex and will be discussed in a subsequent manuscript.

Obtained chromatograms were sorted and analysed using OpenChrom v. 1.3.0 Community Edition. Settings allowing for clean visualisation and peak estimation for piperidine alkaloids were obtained by tracking the ion 98 m/z. This fragment is characteristic of the piperidine moiety of solenopsins, thus allows for better isolation of solenopsins peaks within complex, oft contaminated, field-obtained samples. Different piperidine and piperideine alkaloids were identified from their relative Retention Times (RT) and characteristic ions, as described in previous works (Fox et al.2012, Chen et al. 2009).

## Results

The analyses included 14 fire ant species (both nominal and cryptic species) as listed from Table 1, along with their country of collection. Table 2 depicts the diversity of piperidinic alkaloids retrieved from fire ants from a tentative identification of a total of 32 different compounds observed from the species listed in Table 1. Compounds marked with an asterisk were selected for further use as a reference for species-specific proportions, from being the most predominant and widespread.

**Table 1:** Fire ant species analysed for chemical characters in the present investigation, along with countries of origin. Samples of minor workers were immersed in organic solvents.

| Fire ant species | Country of collection |
| --- | --- |
| *Solenopsis invicta* | Brazil, Argentina, USA, China |
| *Solenopsis richteri* | Brazil, USA, Uruguay |
| *Solenopsis saevissima* cryptic 'Central Atlantic' | Brazil |
| *Solenopsis saevissima* cryptic 'Amazon' | Brazil, Colombia |
| *Solenopsis saevissima* cryptic 'Southern' | Brazil |
| *Solenopsis saevissima* cryptic 'Guyana' | Guadaloupe, French Guyana |
| *Solenopsis saevissima* cryptic 'South Atlantic' | Brazil |
| *Solenopsis metallica* | Brazil |
| *Solenopsis geminata* 'red form' | Brazil, China, USA, French Guyana |
| *Solenopsis geminata* 'dark form' | Brazil, French Guyana, Guadaloupe |
| *Solenopsis virulens* | Brazil, French Guyana |
| *Solenopsis pusillignis* | Brazil |
| *Solenopsis megergates* | Brazil |
| *Solenopsis macdonaghi* | Brazil, Uruguay |

note: a suspected hybrid *S. saevissima* cryptic 'Central Atlantic' x *S. geminata* 'red form' from Brazil is also discussed.

**Table 2:**
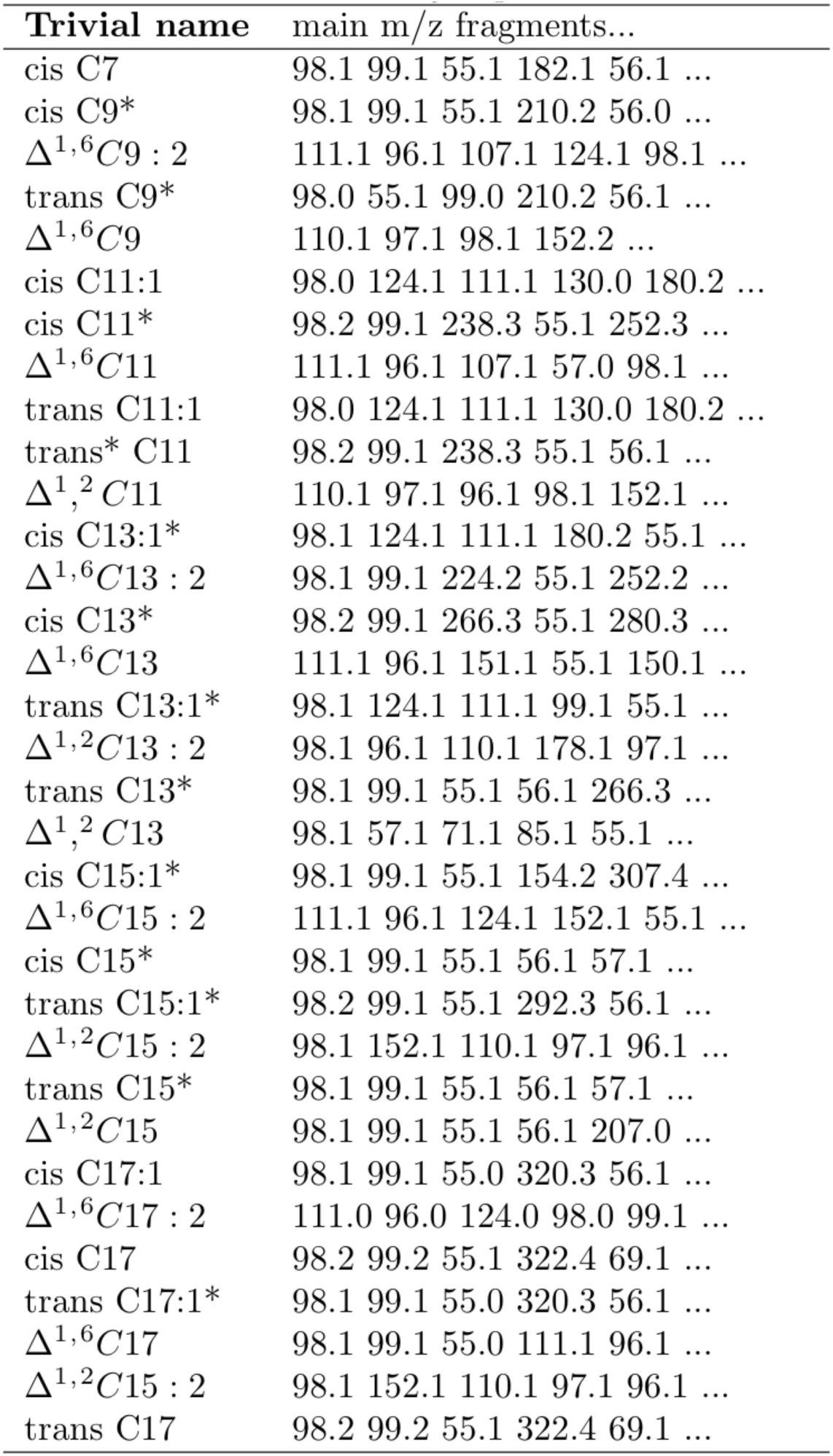
List of piperidine alkaloids assigned to distinct peaks obtained by gas-chromatography analysis of organic solvent samples of fire ants from the New World. Trivial names derive from tentative identifications based on previous studies; m/z fragments by order of magnitude of the top five.

Figure 1 presents the relative proportions of compounds selected from Table 2 as obtained from representative chromatograms of minor workers from the analysed species, therein mainly focusing on C13-rich species of the Solenopsis saevissima species complex. Figure 2 presents the relative proportions of compounds selected from Table 2 as obtained from representative chromatograms of minor workers from the analysed species in Table 1, also including a suspected hybrid, therein mainly focusing on C9-rich species *Solenopsis virulens* and *S. pusignilli*s, and C11-rich species. The obtained proportions proved stable across different samples (full chromatograms to be made available subsequently, along with intraspecific variation), and double-checked with cuticular hydrocarbons patterns whenever possible (hydrocarbons to be discussed in a subsequent manuscript).

**Figure 1.**
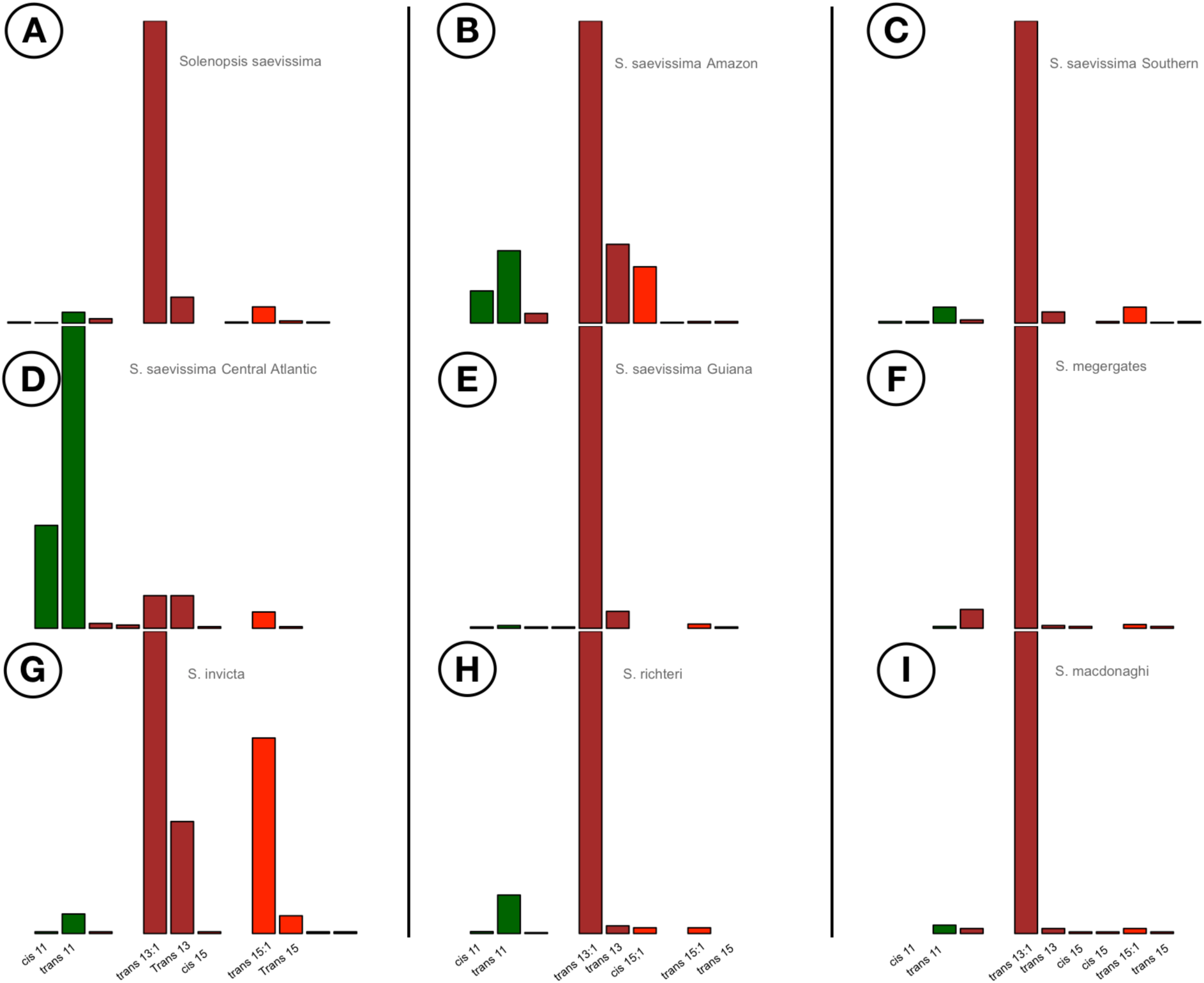
Representative relative proportions of selected venom alkaloids recorded from collected minor workers of different fire ant species — nominal and cryptic species included. Bars represent estimated relative amounts of each compound as estimated by peak area from original selected chromatograms, tracking for the characteristic 98 m/z ion fragment. Colours indicate different side chain lengths, to facilitate visualisation: dark green = 11 carbons; brown = 13 carbons; red = 15 carbons. See Table 1 and references for details on solenopsins structures.

**Figure 2.**
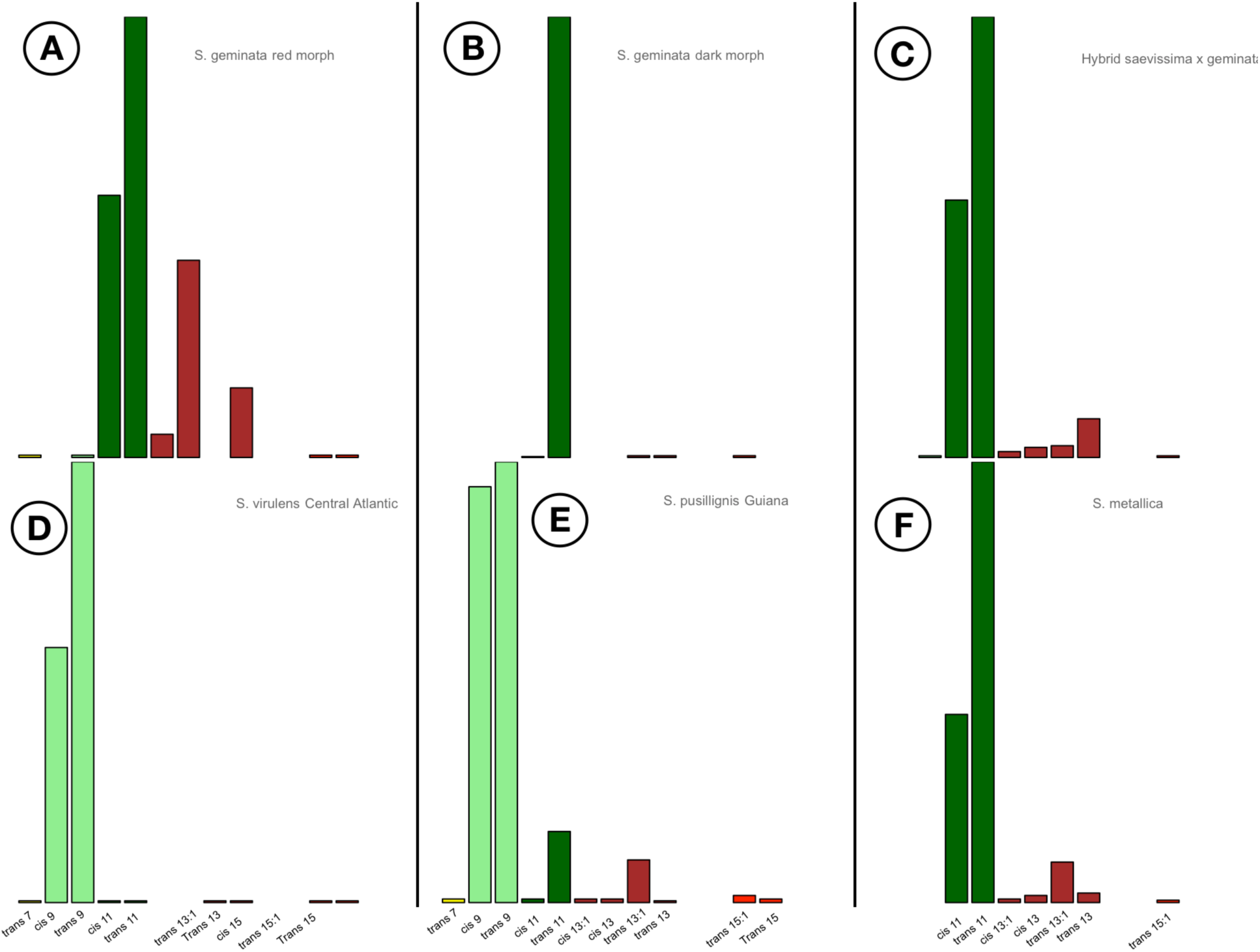
Representative relative proportions of selected venom alkaloids recorded from collected minor workers of different fire ant species — nominal and cryptic species included. Bars represent estimated relative amounts of each compound as estimated by peak area from original selected chromatograms, tracking for the characteristic 98 m/z ion fragment. Colours indicate different side chain lengths, to facilitate visualisation: light green = 9 carbons; dark green = 11 carbons; brown = 13 carbons; red = 15 carbons. See Table 1 and references for details on solenopsins structures.

## Discussion

The provided proportions patterns of piperidine solenopsins among fire ant species are herein given as a proposed additional tool for identifying samples of fire ant minor workers collected from the field into solvents (e.g. ethanol, hexane). As minor workers are usually lacking in clearly defined morphological characters allowing for species identification (Pitts et al. 2005), they are a major challenge for taxonomy.

Piperidine alkaloids are stable and can be recovered from the samples even after a long time elapsed from collection. Tracking for the characteristic 98 m/z ion enables identification of diagnostic peaks amidst dirty and contaminated chromatograms; tracking further compound-specific ions listed in Table 1 provided additional evidence for peak identity. Whilst retention times across variable conditions in different equipments may vary, the order of appearance of the venom alkaloids in chromatograms has consistently remained as presented on Table 1.

Other fire ant species not included in the present study were discussed elsewhere (e.g. *Solenopsis aurea* Wheeler, in MacConnell et al. [1976]), and were also reported to have compounds listed in Table 1. Therefore Table 1 likely lists the total range of piperidinic alkaloids found across *Solenopsis* fire ants, which will be confirmed with of additional analyses from other species.

Earlier studies from the XX century (MacConnell et al. 1976) had often not observed some of the listed compounds in venom chromatograms from several species. Perhaps earlier chromatographs were not sensitive enough to detect compounds at trace amounts. Based on this limitation an evolutionary trend in fire ant venom alkaloids was hypothesised, proposing more derived species such as *S. invicta* and *S. richteri* had evolved more toxic and complex venoms (Brand et al. 1973), as opposed to species within the complex *Solenopsis geminata* which would be biochemically unable to “synthesise piperidines with longer and unsaturated sidechains” (MacConnell et al. 1976, p. 77). It is worth highlighting that species abundant in lower molecular weight solenopsins actually also present trace amounts of the higher molecular weight analogues (e.g. see cis-C15 in *S. virulens*), as demonstrated more recently with *S. geminata* workers and gynes (Fox et al. 2018c). These facts associated with the fact that a number of species within *S. saevissima* can produce the same range of venom alkaloids, in isomeric proportions akin to *S. invicta* and *S. richteri*, weighs against the proposed evolutionary trend. The new evidence indicates the different species must have an equivalent biochemical capacity for producing most (if not all) of the listed compounds.

Therefore the final proportions of venom alkaloids observed in the different fire ant species (compare Fig. 1 and Fig. 2) must be actually regulated by differential expression of key enzymes involved in the final modifications of alkaloid analogues, resulting in differential patterns of the same chemical diversity observed for different species. Deeper understanding of the biochemical pathways and their regulations underlying the production and final modifications of fire ant venom alkaloids will bring more light into the matter.

Regarding general patterns, the most abundant venom alkaloid observed amongst minor workers of different fire ants was *trans* C13:1 (see Fig. 1), followed by *trans* C11 (mainly see Fig. 2). These may well be the most common venom alkaloids in fire ants, overall. It is known that fire ant workers — mainly minor workers—, across all analysed species to date present predominantly trans alkaloid isomers over other analogues (e.g. see Fox et al. 2012). It has been hypothesised that trans alkaloid isomers would the most toxic of such compounds when injected (Greenberg 2008), thus justifying an evolutionary pressure over fire ant workers as active hunters of prey and aggressive nest defendants. However the relative toxicity of the different venom solenopsins is however a controversial matter as the relative toxicity against different model species is bound to vary considerably, also depending on context of application (Fox et al. 2018c). Considering the fact that initial minim workers in early colonies of *S. invicta* abound in C13 analogues (Tschinkel 2006), perhaps the widespread predominance patterns are merely reflecting original precursor pathways and synthesis trail.

Two of the analysed species were abundant in solenopsins C9 analogues, which might be more characteristic of fire ants more closely related with various monomorphic *Solenopsis* thief-ants. Another yet unidentified species reported to about in C9 alkaloids was “Corumba MT Sample #9” reported in MacConnell et al (1974), which was possibly another sample of the *S. pusignilis* population reported herein. Whether such compounds are shared with thief ants will be determined by ongoing investigation into the venom alkaloids of thief ants, and their into the relative abundance. Still, it should be again emphasised not only C9 analogues are observed in the venoms of supposedly more derived species like *S. richteri* (MacConnell et al 1974), and also the presence of trace amounts of longer-side chain alkaloids has been observed from whole nest extractions of *S. virulens* venom(author’s unpublished results), meaning solenopsins of all chain lengths seem biochemically viable for distantly-related species.

In fact, whilst the phylogenetic relationships within different fire ants remain unresolved (Pitts et al. 2018), particularly regarding the different cryptic species of *S. saevissima* (Ross et al. 2010), the observed pattern of venom alkaloids indicates limited correlation with fine species phylogeny. For instance, samples of *S. saevissima* ‘Central Atlantic’ and *S. metallica* are remarkable in being abundant in C11 alkaloids, which is a pattern known from tropical fire ants *S. geminata and* others within the Solenopsis geminata species group. *Solenopsis metallica* was recently described in Pitts et al. (2018) as closely related to *S. megergates* and *S. saevissima*, which may have markedly different chemical profiles. Two colour morphs of *S. geminata* are known from the literature, where a larger, red morph seems to be more widespread than a smaller aggressive *S. geminata* black morph. A recent investigation into their cuticular hydrocarbons suggests they might be two different species (Hu et al. 2017), the reason why their alkaloid profiles are herein presented separately — their taxonomic status is a matter pending deeper investigation. Perhaps a perceptible predominance of *trans* C11 in the venom of *S. geminata* ‘black morph’ minor workers may assist in discrimination in case different species are confirmed. Finally, the alkaloid profile of a suspected hybrid from Brazilian ‘Central Atlantic’ region is presented, based on a single sample. The hybridisation event is suspected from a clear chemical and morphological overlap of characteristic characters between two locally sympatric species. Moreover, evidence pointing to saevissima x geminata hibridisation has been mentioned by Ross et al. (2010) for the same region. This sample will be further analysed in subsequent investigations.

In conclusion, it is hoped that the chemical patterns herein presented may help other researchers in sorting between fire ant samples, whenever freshly collected or original solvents are available to be analysed with a GC-MS system under the settings described. Further detailed analyses regarding intraspecific variations and hydrocarbons will be dealt with in following manuscripts.

## Acknowledgments

The following persons kindly provided help and access to facilities with GC-MS systems: (Agilent 7890A) Li Chen, Yijuan Xu, Xiaoqing Wu, FangYan at IOZ/CAS Beijing and RIFARC/SCAU Guangzhou; (Shimadzu GCMS-QP2010) Georgia Atella, Mileane Bursch, Diogo Gama at HU/ UFRJ Rio de Janeiro; (Thermo-Scientific ISQ QD) Celine Stoffel and Laurent Keller at Biophore/ UNIL Lausanne; (Shimadzu GCMS-QP2010) Odair C. Bueno and Sebastiao Zanao at CEIS/ UNESP Rio Claro; (Agilent 5973A) Anita Marsaioli and Adriana Pianaro at IQ/UNICAMP Campinas. The following persons helped in obtaining insect & chemical samples: Ednildo A. Machado, Robert Lloyd, Sandra F. Fox Lloyd, Maria Elena da Rosa G. Paula, Jose S. L. Martins, Alexandre A. L. Martins, Andrea Bachiao, Daniel R. Solis, Eyal Privman, Meng Xu, Lei Wang, Wen-Wei Tang, HuiJu Zhang, Meng Xu, Muyang He, Dong-Qiang-Zeng, Jerome Orivel, Celine Leroy, Diogo Gama, Rafael R. C. Silva, Oksana Riba, Jason Buser, Adriana Pianaro, Carlos Massuretti de Jesus, Orlando Sireno, Fred, Bruno Reis Braga, Dietrich Gotzek, Jacques Delabie, Henrique G. Bastos, Daniela R. P. Fernandes, Silvia Maria S. Gonçalves. Useful suggestions on data presentation and analysis were provided by Li Chen, Daniel R. Solis. EGPF was funded mainly by CFBE Switzerland during data analysis.

